# Motility in multicellular *Chlamydomonas reinhardtii* (Chlamydomonadales, Chlorophyceae)

**DOI:** 10.1101/201194

**Authors:** Margrethe Boyd, Fran Rosenzweig, Matthew Herron

**Affiliations:** Division of Biological Sciences, University of Montana, Missoula, Montana, USA; Present affiliation: Department of Biomedical Engineering, Northwestern University, Evanston, IL, USA; Present affiliation: School of Biological Sciences, Georgia Institute of Technology, Atlanta, Georgia, USA

**Keywords:** Major evolutionary transition, Volvocales, unicellularity, multicellularity, phototaxis, flagella, trade-off

## Abstract

The advent of multicellularity was a watershed event in the history of life, yet the transition from unicellularity to multicellularity is not well understood. Multicellularity opens up opportunities for innovations in intercellular communication, cooperation, and specialization, which can provide selective advantages under certain ecological conditions. The unicellular alga *Chlamydomonas reinhardtii* has never had a multicellular ancestor yet it is closely related to the volvocine algae, a clade containing taxa that range from simple unicells to large, specialized multicellular colonies. Simple multicellular structures have been observed to evolve in *C. reinhardtii* in response to predation or to settling rate-based selection. Structures formed in response to predation consist of individual cells grouped within a shared transparent extracellular matrix. Evolved isolates form such structures obligately under culture conditions in which their wild type ancestors do not, indicating that newly-evolved multicellularity is heritable. *C. reinhardtii* is capable of photosynthesis, and possesses an eyespot and two flagella with which it moves towards or away from light in order to optimize input of radiant energy. Motility contributes to *C. reinhardtii* fitness because it allows cells or colonies to achieve this optimum. Utilizing phototaxis to assay motility, we determined that newly evolved multicellular strains do not exhibit significant directional movement, even though the flagellae of their constituent unicells are present and active. In *C. reinhardtii* the first steps towards multicellularity in response to predation appear to result in a trade-off between motility and differential survivorship, a trade-off that must overcome by further genetic change to ensure the long-term success of the new multicellular organism.

## Introduction

The evolutionary transition from unicellularity to multicellularity represents a major increase in complexity during the history of life. Higher cell number opens the door to cellular differentiation, enabling diverse processes to be spatially and temporally segregated and to be made more efficient via cellular division of labor. In the volvocine algae, size correlates with level of cellular differentiation. Large-colony members of this clade always contain both somatic and reproductive cells, while small-colony taxa are typically composed of undifferentiated cells that perform both functions. In large colonies, somatic cells are specialized for colony movement and energy production, whereas germ line cells are well-provisioned with resources and dedicated to colony reproduction; this arrangement potentially increases both metabolic efficiency and reproductive potential of the colony as a whole (1). The unicell- to multicell-transition is not an easy one to make, however, with complex life often requiring specific structural and physiological adaptations to overcome a variety of challenges associated with increased size. For example, the diminished ratio of external surface area to volume in a large versus a small organism puts a premium on effective nutrient transport and metabolic waste removal. Increased size associated with the transition to multicellularity is therefore subject to biophysical constraints that can result in trade-offs. These trade-offs may be so severe that the evolutionary success of the new multicellular organism depends on further innovation.

While an integral part of life’s history, the transition from unicellular to multicellular organisms is not yet well understood. Phylogenetic analyses indicate that multicellularity has arisen multiple times in at least 25 independent lineages (2). However, the comparative approach affords limited insight into the genetic changes that led to the first steps along the path towards multicellularity. The comparative approach uses present instead of ancestral relatives due to limited knowledge of ancestral populations, and requires multiple assumptions, including the assumption of a simple and repeatable evolutionary history. To expand our view and to diminish the need for such assumptions, model systems are needed that enable us to simulate the evolution of multicellularity in the laboratory. By experimentally studying this innovation, considered one of life’s major transitions (3), we can gain insight into the evolutionary processes and environmental constraints that may have led to the advent of complex, multicellular organisms.

Given their morphological diversity and the varying levels of differentiation observed among closely related species, the volvocine algae provide an especially useful model system to study the evolution of complexity and multicellularity. *Chlamydomonas reinhardtii*, a unicellular, photosynthetic green alga in the Chlamydomonadaceae, has never had a multicellular ancestor yet is closely related to the volvocine algae, which express multicellularity in colonies of up to 50,000 cells (4). Ranging from simple grouped multicellular clusters to large, differentiated spherical colonies, the volvocine algae show many ways in which multicellularity can be organized. The more complex volvocine algae such as *Volvox carteri* exhibit elaborate structures in which individual cells are anchored and oriented via a shared extra-cellular matrix that also serves as a major interface between cells and their environment. This higher level of organization distinguishes volvocine evolution from the much simpler tetrasporine evolution, in which haploid, thin-walled, nonmotile spores are formed during reproduction (5,6). Species of volvocine algae contain organized groups of *C. reinhardtii*-like cells with varying levels of complexity, indicating that a simple colonial structure composed of unicellular *C. reinhardtii* may be possible.

Predation is thought to favor multicellularity by causing an increased survival rate in clusters too large to be ingested, generating a selective advantage for multicellular groups of algae (7,8). In response to settling rate-based selection, Ratcliff and colleagues have experimentally evolved simple multicellular structures in *C. reinhardtii* (9), while Herron and colleagues have experimentally evolved multicellular structures in response to predation (Person. commun.). Significantly, these *de novo* multicellular lineages continue to form clusters obligately in cultures where no centrifuge or no predator has been present for multiple generations. Two distinct cluster types have arisen in response to selection for large size: clumped, irregular clusters of unicells arose in response to centrifugation, while clusters that evolved in response to predators consist of cells that remain within the parental cell wall, forming clusters as a result of their failure to separate after division. Structure in the latter appears more orderly, as multicell number is frequently a power of two (2^1^ – 2^4^). Because many of these structures are reminiscent of extant volvocine species, the predation isolates were chosen for the present study.

Like all other Volvocales, *C. reinhardtii* is a freshwater organism that is capable of photosynthesis for energy production. *C. reinhardtii* are negatively buoyant, and sink if no effort is expended to maintain their position in the water column (10). Because maintaining an optimal light environment is crucial for its success, it is perhaps unsurprising that *C. reinhardtii* is flagellated and capable of taxis in response to a number of different stimuli. Importantly, the species is capable of phototaxis in response to changes in light intensity. Equipped with two flagella and a single eyespot, *C. reinhardtii* detect differing light levels and use a coordinated breast stroke-like movement to change their position relative to optical stimulus (11). While *C. reinhardtii* is heterotrophic and capable of utilizing environmentally sourced carbon, photosynthesis offers not only additional energy production with carbon uptake, but also an alternative source if environmental carbon is limited (12). As has been demonstrated with *Chlamydomonas humicola*, a close relative of *C. reinhardtii* in the Chlamydomonas genus, mixotrophic cultures have a growth rate about 2.13 times higher than heterotrophic cultures and a rate 7.9 times higher than autotrophic cultures (13). The ability to utilize multiple carbon sourcing pathways has clear implications in the growth rates of algal cultures. Without the ability to undergo phototaxis, *C. reinhardtii* in nature would almost certainly have a diminished capacity to generate energy. Motility is therefore likely to be an important component of fitness in *C. reinhardtii*.

Multicellular relatives of *C. reinhardtii* such as *Gonium, Eudorina* and *Volvox* also rely on flagellar movement. Volvocine algae have been observed to have exteriorly oriented flagella regardless of colony complexity. These negatively buoyant multicellular species rely on flagellar movement not only to respond to changes in light environment but also to increase nutrient flow around the colony, facilitating nutrient transport (10).

Here we explore the capacity of *de novo* multicellular strains of *C. reinhardtii* to undergo phototaxis. Our goal is to evaluate the impact of the first steps in a major transition on motility, a determinant of fitness in this freshwater phototroph. By evaluating the capacity of multicellular *C. reinhardtii* to move in response to a key environmental stimulus, we can gauge its potential viability in nature and identify what further evolutionary innovations – and selection pressures – would be needed to evolve differentiated multicellularity in the laboratory.

## Methods

*Strain construction and culture conditions*. Experimental populations of *C. reinhardtii* were generated by crossing parental strains CC1690 and CC2290, resulting in a genetically diverse pool of F2 progeny. Subsamples of this outcrossed population were then used for all treatment and control groups. *C. reinhardtii* in the treatment groups were co-cultured with the predator *Paramecium tetraurelia*, while those in the control groups were cultured axenically. Algae were cultured in 24-well plates using WC media (14), with the experiment running for *ca*. 300 generations over six months. Eight strains were isolated at random from each population through three rounds of serial plating, then cultured in 3 mL TAP media (15) and stored on TAP + 1.5% agar plates. Strains were regrown in 5 mL WC at a temperature of 22.5°C under cool white illumination at 4100 K. Cultures provided upon request.

*Assay of motility in experimentally evolved algae*. Motility in response to light was evaluated on cell lines that displayed varying degrees of multicellularity, as well as a unicellular control population. Three populations of *C. reinhardtii* were grown until dense enough to see with the naked eye, a density of about 2 ×10^5^ cells/mL. Eight strains were isolated from each of two experimental populations and one control population. The control population, K1, contained eight unicellular strains, experimental population B2 contained three unicellular strains and five multicellular strains, and experimental population B5 contained eight multicellular strains (**Fig. 1**). Once grown ~2 ×10^5^ cells/mL, cultures were diluted to the same absorbance of 0.1 using a spectrophotometer set at λ=800 nm. The 24-well tissue culture plates used had their longitudinal textured edges removed to reduce light scattering and to expose wells directly to the light source. Strains were randomized in such a way as to ensure one replicate of each strain per plate, at least one multicellular and one unicellular strain per plate, and varying replicate position between plates to reduce positioning effects. Using this randomized order, strains were placed into the six wells along the long side of the modified plate to ensure as equal light distribution among wells as possible.

**Figure 1.**
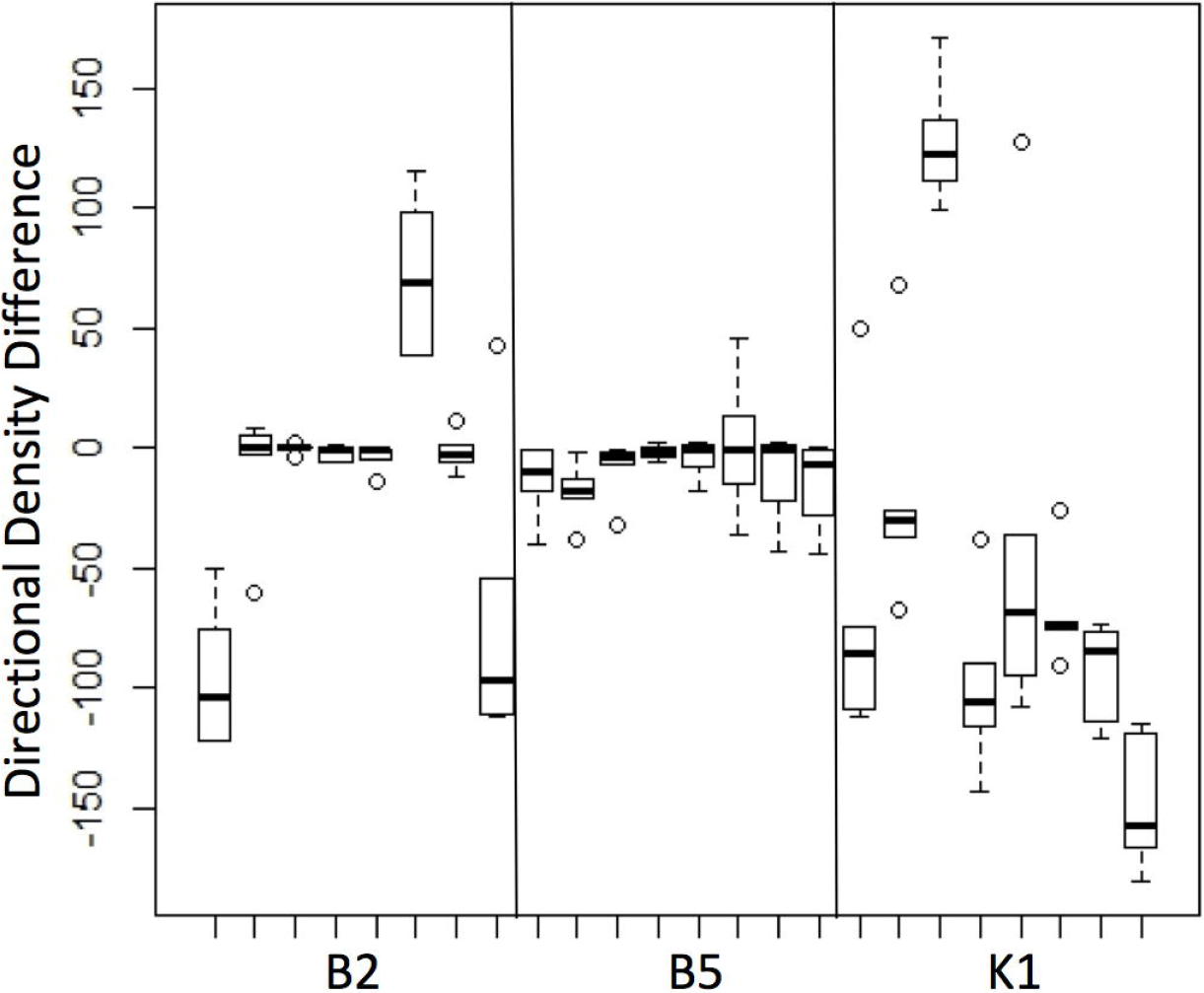
Multicellular and unicellular phenotypes. Cell populations from the B2 population (a), K1 population (b) and B5 population (c). (images by Josh Ming Borin)

To measure phototaxis, a copy stand, designed to hold camera positioning and light level steady, was mounted with a Canon Rebel T3 Powershot camera equipped with a Canon EF-S 18-55mm f/3.5-5.6 IS II lens. Modified tissue culture plates were placed on the platform of the copy stand with the algal wells facing a cool white (4100 K) directional incubator light. Light level at the plate was fixed at 170 lux. An image of the plate was taken before the directional light was switched on using the neutral light provided by the copy stand and a cable release to minimize camera movement. Then, the copy stand lights were turned off, the directional light was turned on, and plates were exposed from one edge for a period of five minutes. Light from all other sides was blocked. Once the five-minute period had elapsed, the directional light was switched off, copy stand lights switched on, and a final image was taken. Because cultures were grown to a density that could be seen with the naked eye, density changes in the well due to algal phototaxis were immediately visible during and after light exposure. The images taken recorded these visual changes for later analysis. This process was repeated with all plates; ultimately six sets of plates were analyzed and six replicates per strain were obtained (**Fig. 2**). Each replicate was assayed one time, and separate cultures were used.

**Figure 2.**
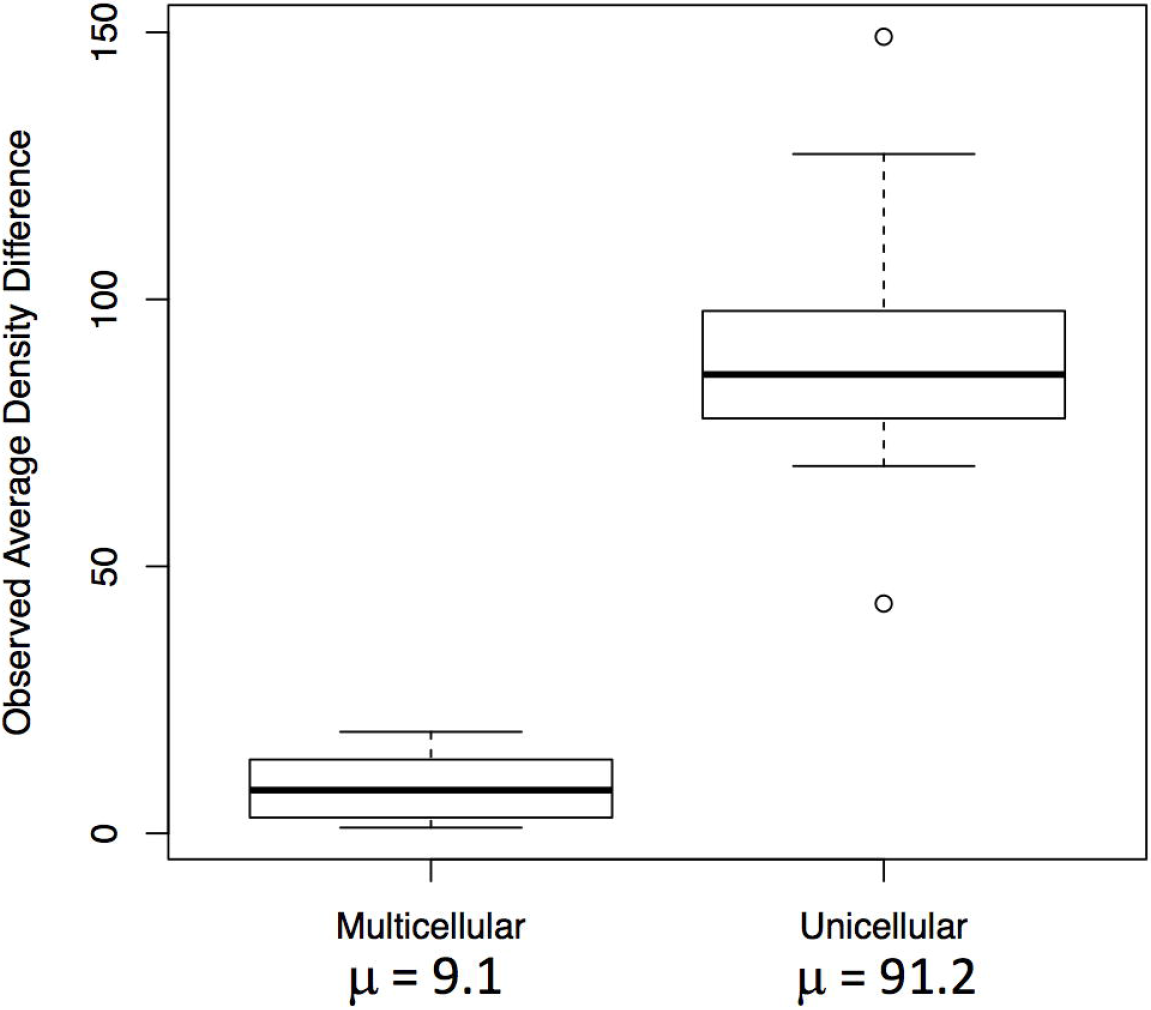
Experimental Design. Illustration of the experimental design of motility assays.

To compare levels of phototaxis between unicellular and multicellular strains, images were loaded into *ImageJ* photo-analysis software for analysis (16). For each set of “before” and “after” images, density changes were analyzed by comparing the pixel density changes over the time-course of light exposure. Because camera position and lighting were held constant between the images, changes in pixel density could be attributed to algal cell movement within the well in response to light. In order to determine net change in pixel density, we subtracted one image from the other. This procedure creates a map of change in algal density over the five-minute exposure time. To further improve our analysis, the color threshold on this map could be set to display only green-spectrum pixel changes. This helps to reduce any possible effects due to shadowing or slight movements of the plate or camera, and specifies our analysis for green cells and clusters of *C. reinhardtii*. For each well in each plate a plot profile was obtained to determine where the highest density changes of algae occurred and in which strains. The difference between the average density of the lighter half of the well (closer to the directional light source) and that of the darker half was then taken to obtain the direction of movement, and to indicate no directional movement if this difference proved insignificant. In this way, a relative magnitude and direction of phototaxis was measured in each strain (**Fig. 3**, **4**).

*Microscopy*. To further explore why movement is limited in multicellular *C. reinhardtii*, strains were observed under a light microscope to determine whether structural features peculiar to multicellular colonies might limit motility. To preserve colonial integrity and viability no stains or fixing agents were used. Microscopy was done at 80x magnification.

**Figure 3.**
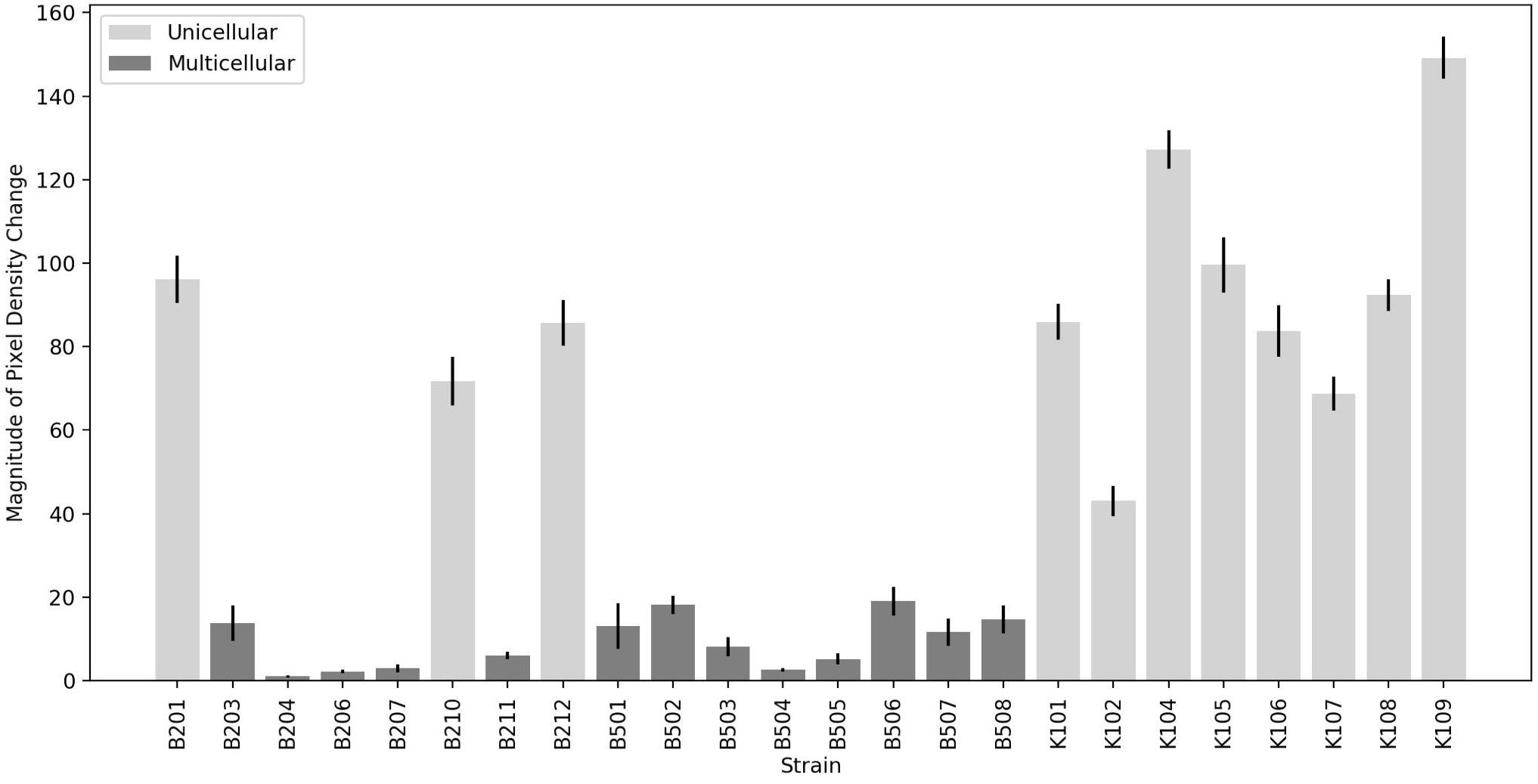
Well Plate Before and After Light Exposure. Longitudinal wells in a modified 24-well plate before and after light exposure. Image A shows the plate before exposure, while image B was taken after light exposure. Black arrows indicate changes in algal density due to directional light exposure. Scale bar = 8mm.

**Figure 4.**
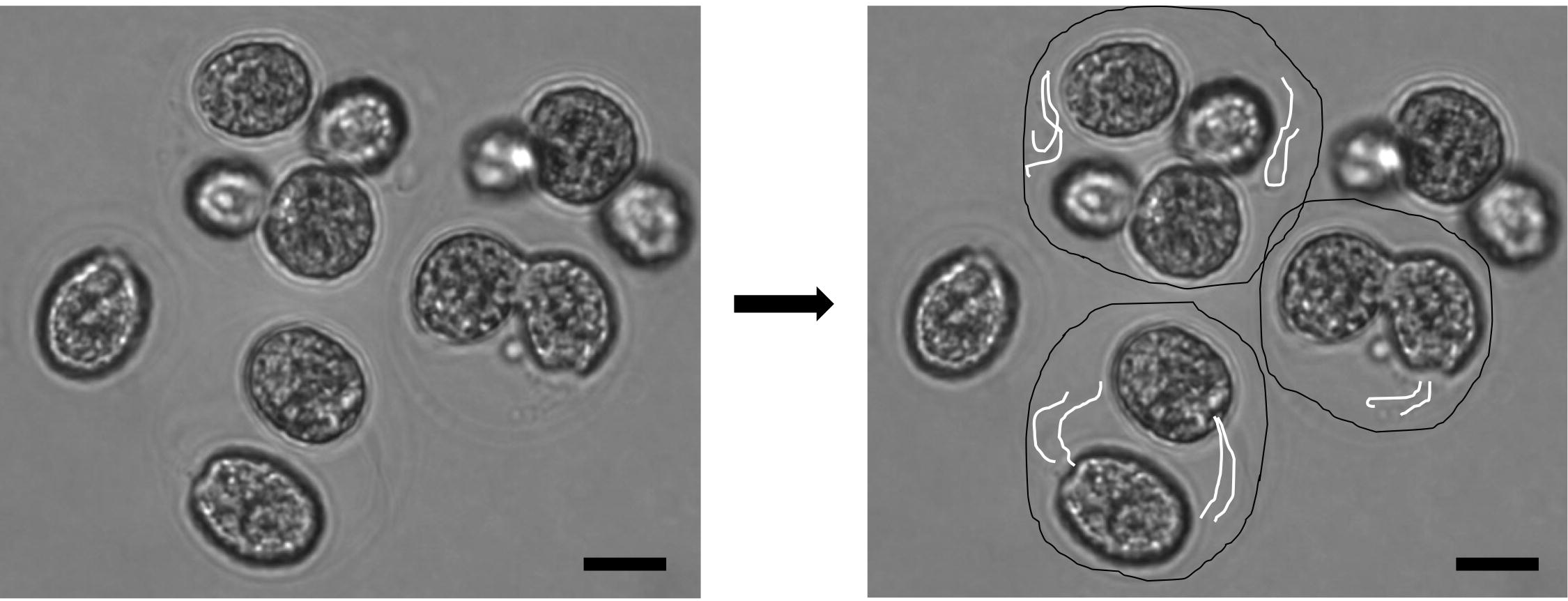
Well Plate Analysis in *ImageJ*. Images from Figure 2 analyzed to indicate density differences in the plate before and after directional light exposure. Image A shows the *ImageJ* analysis showing the changes in density of green pixels between the before and after photographs, showing algal movement due to light exposure. Image B shows this same analysis image with a color threshold set to determine relative magnitude of algal movement. Scale bar = 8mm.

## Results

Microscopy indicates that in the multicellular form, unicellular flagellated *C. reinhardtii* appear to be encased within an external wall with few protrusions into the surrounding media. (**Fig. 5**). We hypothesize that in the case of predation-selected lines, multicellularity evolved by failure of daughter cells to separate after cell division. If so, then each colony consists of a group of daughter cells trapped within the maternal cell wall. Importantly, each individual algal cell within the confines of the group remains flagellated and can be observed moving these flagella within the colony (see supplemental video **Fig. S1**). In some cases, a few flagella were observed to be moving outside of the colony wall. Phototactic assays were performed to determine whether these flagella and those inside the colony can generate movement despite being largely confined within the nascent colony.

**Figure 5.**
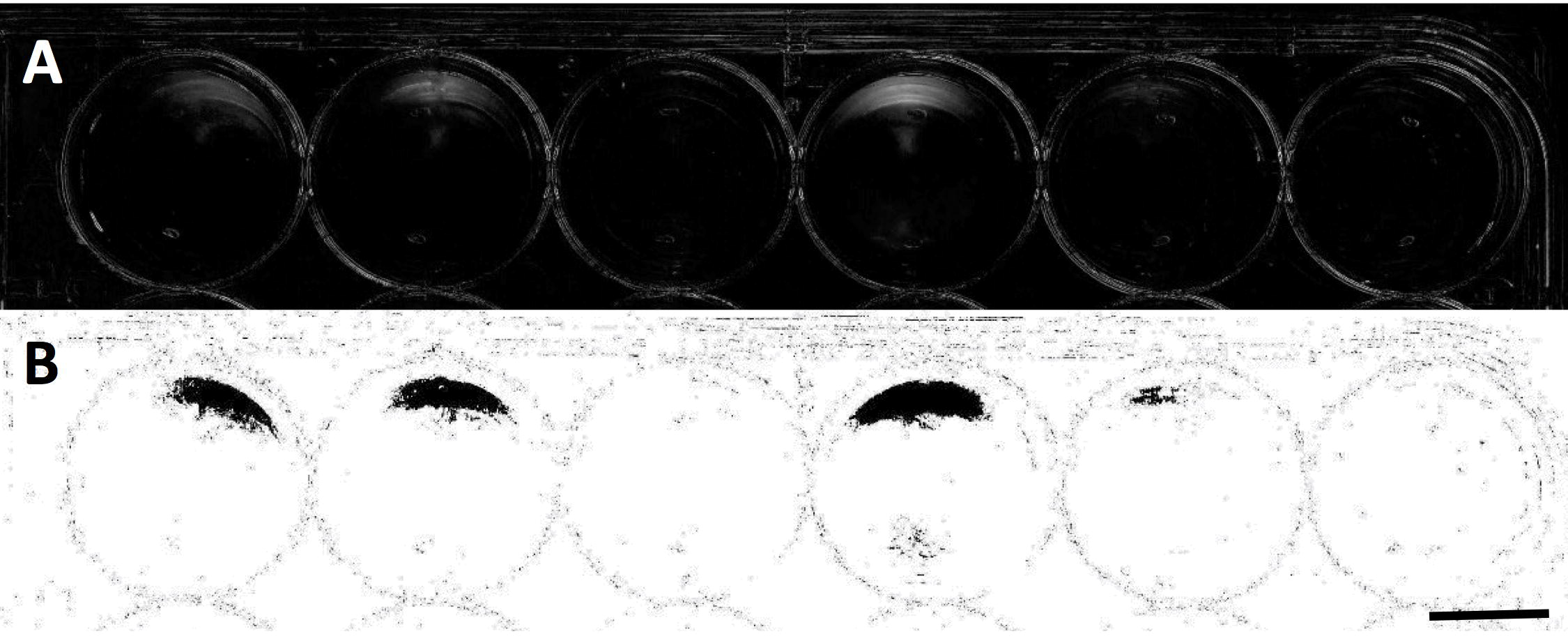
Microscopy of Multicellular Colony. Multicellular colonies appear to remain within parental cell wall and retain visible flagella, which were observed to be capable of movement. In the right-hand version of this image black lines have been drawn to better indicate potential colony walls, while white lines indicate flagella. Scale bar = 10μm

**Figure 6.**
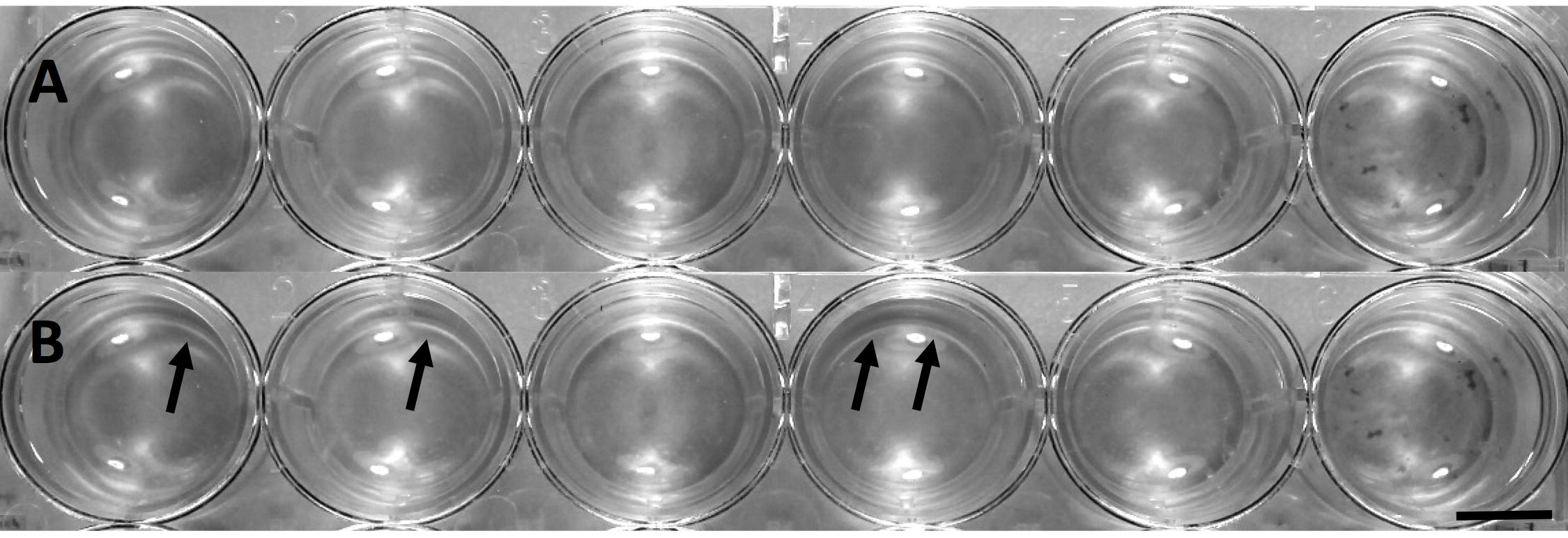
Density Changes per Strain. Change in algal density distributions in unicellular and multicellular strains due to light exposure.

In phototactic assays multicellular strains of *C. reinhardtii* exhibited very limited – if any – phototaxis (**Fig. 6**). Because direct microscopy indicates that the multicellular stages are largely immobile (see supplemental video **Fig. S1**), we hypothesize that the low levels of phototaxis observed in multicellular strains (**Fig. 6, 7, 8**) were due to the presence of two phenotypes in these cultures: multicellular colonies and motile unicellular propagules released during the colonial reproductive cycle.

Transmission microscopy revealed that multicellular cultures contain a small number of these motile unicellular propagules, which are released during multicellular strains’ reproductive cycle when a colony reaches its characteristic burst size. These propagules remain phototactic, indicating that the flagellar function and phototactic ability of individual algal cells are not impaired. Importantly, these unicellular propagules obligately develop into multicellular colonies after their release from the parent colony, even in the absence of a predator. It is this non-plastic, obligate colony formation in particular that distinguishes these strains from palmelloid colonies, the formation of which has also been observed in *C. reinhardtii* in the presence of certain environmental factors (9,17). Some strains, when viewed under a microscope at 80x magnification, had a small number of these unicellular propagules swimming among the multicellular clusters regardless of when strains were analyzed, so a slight response is expected due to these motile propagules.

**Figure 7.**
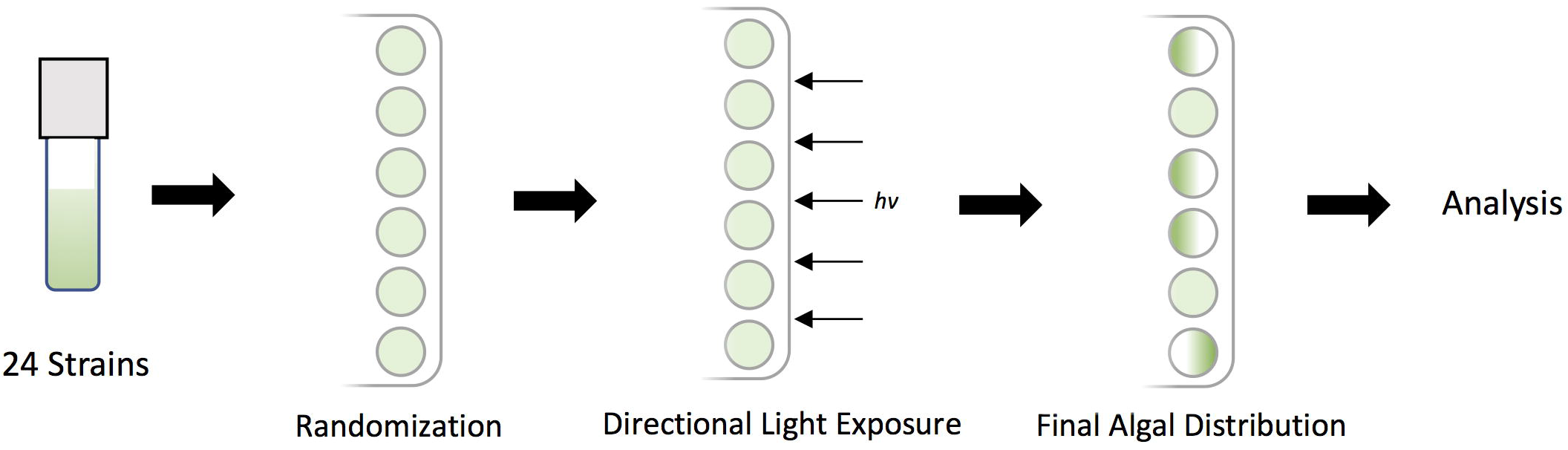
Magnitude of Phototaxis. Analysis of the relative magnitude of algal movement due to light exposure between multicellular and unicellular strains.

**Figure 8.**
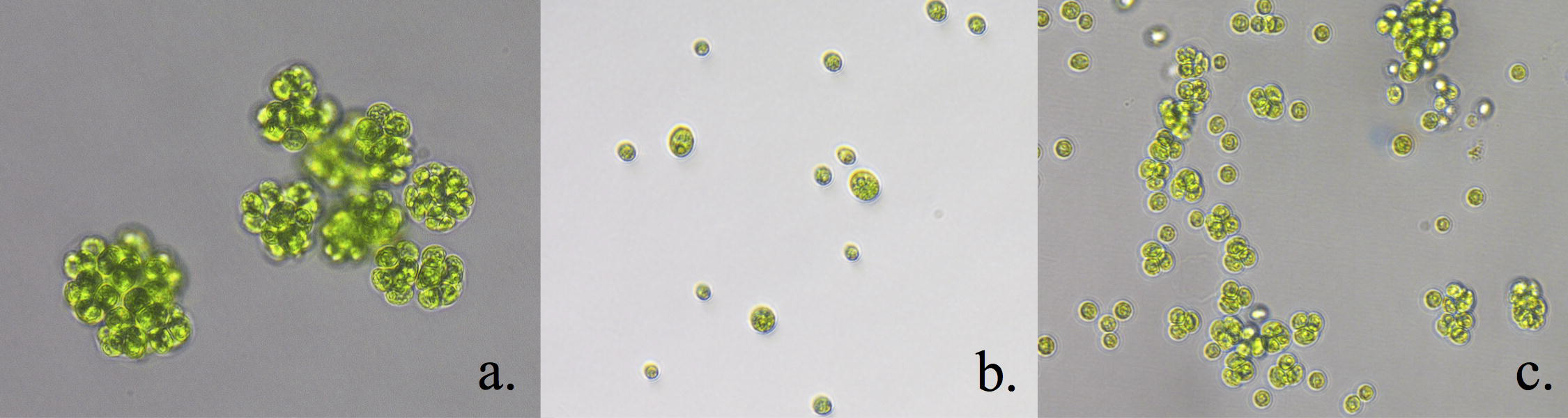
Directionality of Phototaxis. Analysis of algal movement due to light exposure where positive values indicate accumulation in the half of the well toward the light source and negative values indicate accumulation in the other half.

Despite this small margin of error, levels of phototaxis in unicellular strains were ten times higher on average than levels of phototaxis in multicellular strains. Thus, cells within multicellular units that evolve from unicellular ancestors appear to lose the capacity to be motile when in a colony, at least in the earliest stages of this transition. While capable of individual movement within the colony, cells can neither move themselves out of the colony nor move the colony as a whole because too few flagella extend past the cell wall to exert force against the surrounding fluid. From this, we can conclude that the advent of multicellularity in *C. reinhardtii* severely limits motility in comparison to unicellular ancestors.

All unicellular strains observed in this experiment exhibited significant directional movement. The direction and magnitude of movement differed among strains, though this is not necessarily an unusual result when analyzing genetically diverse populations as direction and level of phototaxis has been shown to be affected by a number of factors; these include the presence of reactive oxygen species, varying CO_2_ concentrations, and other environmental factors (11,18). Two strains exhibited strong negative phototaxis at the same light intensity that all other strains exhibited positive phototaxis; moreover, among unicells values for positive phototaxis varied over an order of magnitude (**Fig. 7**). The two negatively phototactic strains came from different populations: one came from the B2 experimental population, the other from the K1 control population. These results indicate differential light sensitivity in unicellular strains. Despite these varying phototactic responses, however, all genetically unicellular strains were phototactic overall. This indicates a significant difference between varied response in unicellular strains and immobility in multicellular strains, suggesting that the absence of motility in multicellular strains is not attributable to an environmental cause or simply to differential phototactic response. Despite variations of this sort, however, no flagellar defects or other motility-limiting factors were observed in genetically unicellular strains or in unicellular propagules from multicellular colonies, indicating that the process of selection by predation did not hinder unicellular movement. When average density changes were compared using ANOVA, there was an order of magnitude difference between average density changes of multicellular versus unicellular strains (*p* < 0.0001) (**Fig. 7**).

## Discussion

In multicellular *C. reinhardtii* strains evolved under predation, structure limits motility. With flagella internalized, no amount of beating can alter the position of the colony. In other volvocine species externally oriented flagella are a key characteristic of the algal colony, and elegant mechanisms have evolved to regulate and coordinate their movement (5,10). We see this in species as simple as the regularly clumped *Gonium* to species as complex as *Volvox*, whose colonies contain thousands of cells. The use of these flagella is critical and their purposes are multifaceted, indicating that a loss of function would be highly detrimental. Flagella facilitate boundary layer mixing and nutrient transport as well as motility, and all three functions contribute to algal viability (10,19). In our experiments, populations were evolved in illuminated incubators with ample nutrients obviating the need for photo- or chemotaxis. Thus, while our evolved multicells thrive in a laboratory environment their immobility would likely be detrimental to fitness in nature. Additional or alternative selection pressures will be required to study how motile multicellular colonies might evolve from unicells in natural environments. These selective measures might include the use of a chemical gradient or a light gradient in conjunction with predation, as described below.

Cells within *de novo* multicellular *C. reinhardtii* colonies remain capable of individual flagellar movement and are often able to move themselves within the colony (see supplemental video, **Fig. S1**). In addition, when colonies break apart during their life cycle unicellular propagules remain capable of phototaxis. Thus, de novo multicellular strains appear not to have lost photo-sensing ability or flagellar function under selection for large size. This characteristic differentiates these nonmotile colonies from simple tetrasporic groups, as a shared cell wall is present but cells within the wall remain flagellated and motile (6). We also observe that despite consistent treatment through the experiment, levels and direction of phototaxis vary widely among strains. Cultures serially propagated at the same temperature under identical light and nutrient levels responded very differently to light stimuli. Because some strains were highly phototactic while others were not, it appears that some strains are more responsive to light stimulus than others. This is not an unexpected result, as various factors can affect levels of phototactic response in genetically diverse populations (11,18).

While structural barriers are the most obvious factors hindering motility, the lack of cellular coordination may be a significant hindrance as well. Colonies cannot easily accomplish directional movement if all cells are acting independently. In larger volvocine colonies, cells must synchronize their activities to create a pattern of flagellar movement that can move the group (10). This coordination is made possible through the existence of the highly structured extra-cellular matrix anchoring cells in the correct orientation relative to one another (5). Because we cannot observe free (external) flagellar movement in our multicellular strains we cannot ascertain whether their movement is in any way coordinated, however we observe that any structural organization offered by the external matrix is not present.

In conclusion, experimental laboratory evolution can be used not only to explore metabolic adaptation (20,21) and life history evolution (22), but also to identify the genetic pathways and ecological pressures that promote major evolutionary transitions (9). By examining the motility phenotype of newly evolved multicellular *Chlamydomonas* we have uncovered a trade-off that arises from their novel morphology. However, because flagellar structure and function are unchanged, and unicellular propagules remain phototactic, barriers to the subsequent evolution of externalized flagella and coordinated movement in multicells may not be too high to overcome in the laboratory. Multiple, and possibly sequential, selective pressures are likely required to experimentally evolve naturally viable multicellular algal clusters. Identifying the nature of these selection pressures, as well the order and intensity required to produce differentiated multicellularity promises to yield insight into mechanisms underlying a major evolutionary transition in the history of life on Earth.

## Acknowledgements

Funding for this work was provided by the NASA Astrobiology Institute (Cycle 7 Cooperative Agreement Notice), the National Science Foundation (DEB-1457701) and the John Templeton Foundation (43285).

